# SARS-CoV-2 Viroporins Activate The NLRP3-Inflammasome Via The Mitochondrial Permeability Transition Pore

**DOI:** 10.1101/2022.02.19.481139

**Authors:** Joseph W. Guarnieri, Alessia Angelin, Deborah G. Murdock, Prasanth Portluri, Timothy Lie, Douglas C. Wallace

## Abstract

Cytokine storm precipitated by activation of the host innate immune defenses is a major cause of COVID19 death. To elucidate how SARS-CoV-2 initiates this inflammatory process, we studied viroporin proteins E and Orf3a (2-E+2-3a). Expression of 2-E+2-3a in human 293T cells resulted in increased cytosolic Ca^++^ and then elevated mitochondrial Ca^++^, taken up through the MUCi11-sensitive mitochondrial calcium uniporter (MCU). Increased mitochondrial Ca^++^ resulted in stimulation of mitochondrial reactive oxygen species (mROS) production, which was blocked by mitochondrially-targeted catalase or MnTBAP. To determined how mROS activates the inflammasome, we transformed 293T cells with NLRP3, ASC, pro-caspase-1 and pro-IL-1β plus used THP1 derived macrophages to monitor the secretion of mature IL-1β. This revealed that mROS activates a factor that is released via the NIM811-sensitive mitochondrial permeability pore (mtPTP) to activate the inflammasome. Hence, interventions targeting mROS and the mtPTP may mitigate the severity of COVID19 cytokine storms.

## INTRODUCTION

Approximately 350 million cases of COVID19 have been reported globally, resulting in over 5.5 million deaths (Dong et al., 2020). COVID19 is caused by SARS-CoV-2 whose genome structure encodes a polyprotein that is cleaved into 16 non-structural proteins (nsp) as well as the structural proteins S (Spike), E (Envelope), M (Membrane), and N (Nucleocapsid), and seven open reading frames (orfs) 3a, 6, 7a, 7b, 8, 9b, and 10, with substantial homology with SARS-CoV-1.

Severe COVID19 manifests as pneumonia, acute respiratory distress syndrome, respiratory failure, and cytokine storm resulting in multiple organ failure (Ferreira et al., 2021; Rodrigues et al., 2021; Yang et al., 2021). The cytokine storm results from the elaboration of pro-inflammatory cytokines such as interleukin (IL)-1β (Ajaz et al., 2021; Chen et al., 2020; Chi et al., 2020; Han et al., 2020; Lucas et al., 2020; Wen et al., 2020).

The production of mature IL-1β requires the activation of the mitochondrially-bound NLRP3-inflammasome (NLRP3-I), which encompasses the NLR family pyrin domain containing 3 (NLRP3) receptor; the adaptor molecule apoptosis-associated speck-like protein containing a caspase activation and recruitment domain (ASC); and the pro-IL-1β-converting enzyme pro-caspase-1 (CASP1). Upon activation, the NLRP3-I triggers the proteolytic cleavage of pro-caspase-I (pro-CASP1), and CASP1 cleaves pro-IL-1β to generate IL-1β which is secreted from the cell (Broz and Dixit, 2016). Autopsy samples from severe COVID19 patients display increased NLRP3-I activation in lung tissues and peripheral blood mononuclear cells (Rodrigues et al., 2021), and monocytes isolated from severe COVID19 patients have increased levels of activated NLRP3-I and IL-1β, (Ferreira et al., 2021). Thus, understanding the mechanism by which SARS-CoV-2 activates the NLRP3-I is imperative for understanding the pathophysiology of severe COVID19.

Recently, mitochondrial dysfunction has been shown to activate the innate immune system via mitochondrial reactive oxygen species (ROS) production and oxidation of the mitochondrial DNA (mtDNA) during replication, induced by the expression of the rate-limiting enzyme cytosine monophosphate kinase 2 (CMPK2). The oxidized mtDNA (Ox-mtDNA) is released from the mitochondrion to bind and activate the NLRP3-I (West and Shadel, 2017; Zhong et al., 2018). While the mechanism by which SARS-CoV-2 activates the NLRP3-I is unknown, expression of the SARS-CoV-1/2 viroporins have been associated with activation of NLRP3-I (Chen et al., 2019, Nieto-Torres, 2015 #118; Siu et al., 2019; Xia et al., 2021; Yue et al., 2018) and are known to be membrane ion channels (Hover et al., 2017; Nieva et al., 2012).

SARS-CoV-2 encodes two viroporins E (2-E) (Verdiá-Báguena et al., 2021) and ORF3a (2-3a) (Qu et al., 2021), with homologues to the SARS-CoV-1 proteins (Kern et al., 2021; Mandala et al., 2020). SARS-CoV-1/2 E and 3a viroporins localize to the endoplasmic reticulum (ER), Golgi apparatus, and plasma membrane (Gordon et al., 2020a) where they increase the permeability to cations such as Ca^++^ (Minakshi and Padhan, 2014; Verdiá-Báguena et al., 2021, Kern, 2021 #420; Verdiá-Báguena et al., 2012). For SARS-CoV-1, the 1-E and 1-3a have been shown to activate the NLRP3-I in human monocyte-derived macrophages (Chen et al., 2019; Siu et al., 2019; Yue et al., 2018). In LPS-primed macrophages, co-expression of 1-E plus 1-3a resulted in higher levels of IL-1β secretion than either viroporin alone (Chen et al., 2019), 1-E has been reported to activate NLRP3-I through disrupting Ca^++^ homeostasis in cells (Nieto-Torres et al., 2015; Xia et al., 2021), and activation of the NLRP3-I and secretion of IL-1β by 1-E and 1-3a is mitigated by treatment with the mROS scavenger MitoQ (Chen et al., 2019). However, the mechanism by which SARS-CoV-2 activates the NLRP3-I has yet to be explained.

We hypothesized that expression of 2-E plus 2-3a results in increased Ca^++^ flux into the cytosol where it is taken up by the mitochondrion through the mitochondrial Ca^++^ uniporter (MCU). Within the mitochondrion, the Ca^++^ activates the pyruvate and α-ketoglutarate dehydrogenases to generate excessive NADH (Denton, 2009). The increased NADH overloads the mitochondrial electron transport chain producing increased mitochondrial ROS (mROS). The mROS oxidizes the mtDNA, and the Ox-mtDNA is released through the mitochondrial permeability transition pore (mtPTP) to bind to the NLRP3 inflammasome. This activates caspase-1 to cleave pro-IL-1β resulting in the secretion of active IL-1β (Xian et al., 2021; Zhong et al., 2018). Our current results support this scenario, thus placing mitochondrial function at the nexus between SARS-CoV-2 infection and the cytokine storm of severe COVID19.

## RESULTS

### Expression of 2-E+2-3a increases Ca^*++*^ leakage into the cytosol, elevates mitochondrial Ca^*++*^ levels, and increases mROS production

We constructed a polycistronic expression vector combining 2-E+2-3a (LV-E3a) (**Figure 1A&B**). In this vector the 2-E+2-3a sequences were separated by the self-cleaving 2A peptide site (Liu et al., 2017; Szymczak-Workman et al., 2012) to allow co-expression from a single transcript. The expression of 2-E+2-3a in LV-E3a transduced 293T cells was confirmed by Western blot (**Figure 1C**). We next demonstrated that LV-E3a transduced 293T cells experience increased cytosolic Ca^++^ with Fura-Red (**Figure 1D-F**) and mitochondrial Ca^++^ with Rhod2 (**Figure 1G-I)**. Thapsigargin (TG), which triggers Ca^++^ flux into the cytosol, increased mitochondrial Ca^++^ uptake (Bagur and Hajnóczky, 2017; Csordás et al., 2018), and mitochondrial Ca^++^ uptake was increased in TG treated LV-E3a transduced cells. Co-treatment with the mitochondrial calcium uniporter inhibitor 11 (MCUi11) (Di Marco et al., 2020; Márta et al., 2021) abolished mitochondrial calcium uptake (**Figure 1J**). Thus, expression of 2-E+2-3a in 293T cells results in elevated cytosolic Ca^++^ which is taken up by the mitochondrial calcium uniporter resulting in elevated mitochondrial Ca^++^ (**Figure 1J**).

**Figure 1.**
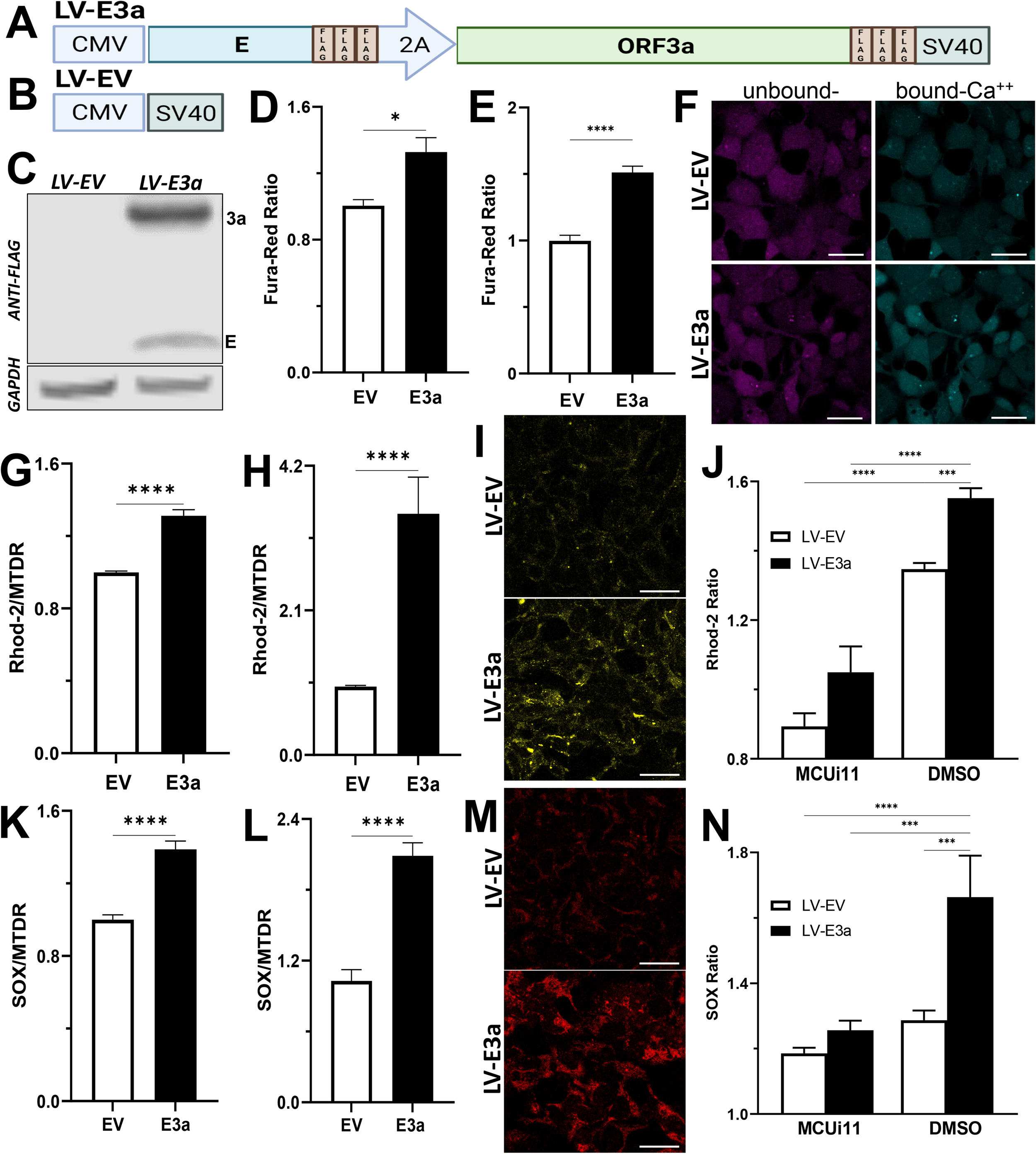
Expression of 2-E+2-3a induces mROS production by elevating mitochondrial Ca^*++*^ levels. **A-B)** Schematics of our LV-EV and LV-E3a vectors. **C)** 293T cells were transduced with LV-EV or LV-E3a, and samples analyzed by immunoblot with a mouse monoclonal antibody against FLAG-Tag, to detect the FLAG-tagged 2-E+2-3a viroporins. GAPDH was a loading control. **D-F)** 24 hrs post-transduction cells were stained with Fura-Red to measure cytosolic Ca^++^ levels, **G-I)** Rhod2 & Mitotracker Deep Red (MTDR) to measure mitochondrial Ca^++^ levels, or **K-M)** MitoSOX and MTDR to measure mROS levels via **E-F. H-I, L-M)** data from confocal microscopy or **D, G, K)** data from plate reader assays. **J, N)** 24 hrs post-transduction with LV-EV or LV-E3a 293T cells were treated with or without 10 μM MCUi11, and then 2.5 μM TG and stained with **J)** Rhod2 to measure mitochondrial Ca^++^ levels, or **N)** MitoSOX to measure mROS levels by plate reader assays. Scale bar = 30 μm. Error bars represent SEM from 3 independent experiments; statistically significant data is indicated with asterisks (*).

We next demonstrated that 2-E+2-3a expression in 293T cells increased mROS production by staining transduced cells with MitoSOX, which detects mitochondrial superoxide anion production. MitoSOX fluorescence confirmed that 2-E+2-3a expression increased mROS production (**Figure 1K-M**). To determine if the increased mROS was due to the entry of Ca^++^ into the mitochondrion, we treated the cells with MCUi11which blocked the increased mROS production (**Figure 1N**).

To confirm that the 2-E+2-3a induced ROS production was mROS, we transformed the 2-E+2-3a expressing 293T cells with a vector expressing mitochondrially-targeted catalase (mCAT) (**Figure 2A-C**) which removes mitochondrial H2O2 (Schriner et al., 2005), or treated the cells with the mitochondrially targeted catalytic metalloporphyrin anti-oxidant, MnTBAP (Melov et al., 1998; Tong et al., 2007). Treatment with either MnTBAP or mCAT extinguished the 2-E+2-3a activated ROS product, confirming that the ROS was generated by the mitochondrion (**Figure 2D-I**).

**Figure 2.**
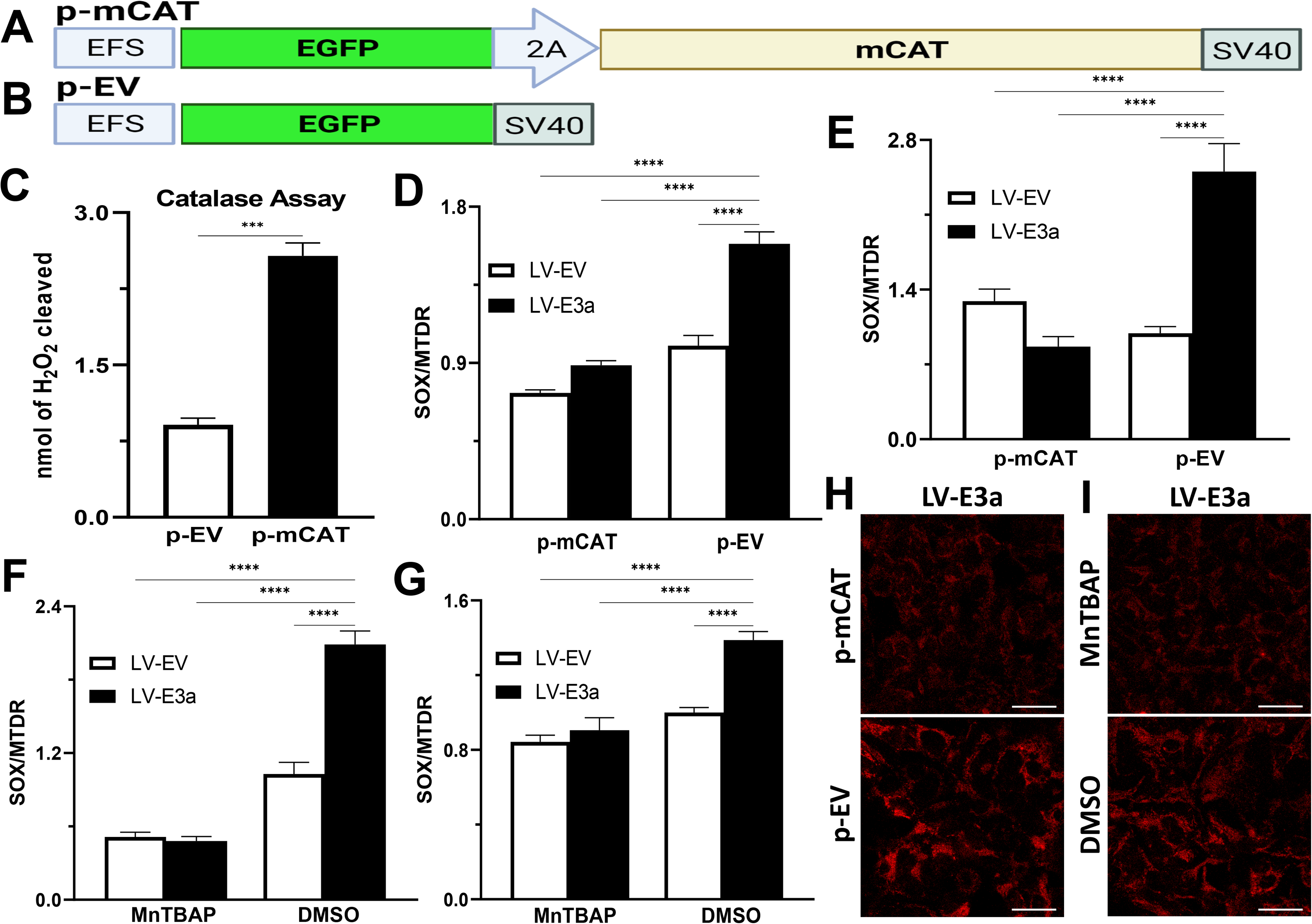
Expression of mCAT or treatment with the mROS scavenger MnTBAP antioxidant defenses blocks 2-E+2-3a induced mROS. **A-B)** Schematics of p-EV and p-mCAT vectors. **C)** Catalase assay through cleavage of H_2_O_2_ in 293T cell lysates collected 24 hrs post-transfection with p-EV or p-mCAT. **D-I)** 24 hrs post-transfection levels of mROS assessed using MitoSOX and MTDR fluorescence 293T cells transduced with LV-EV or LV-E3a and transfected **D, E, and H)** with p-mCAT or its respective control p-EV or **F, G, and I)** cultured in the presence or absence of 50 μM MnTBAP, DMSO used as a negative control. MTDR fluorescence was used to normalize for mitochondrial content with mROS expressed as the ratio of MitoSOX/MTDR, by **D, F)** confocal microscopy or **E, G)** plate reader assays **H, I)** Representative images of MitoSOX-stained cells. Scale bar = 30 μm. Error bars represent SEM from 3 independent experiments; statistically significant data is indicated with asterisks (*).

### 2-E+2-3a induced mROS is involved in NLRP3-activated and IL-1β production

In SARS-CoV-1, activation of the NLRP3-I and pathogenicity are associated with both 1-E+1-3a (Chen et al., 2019; Nieto-Torres et al., 2014; Xia et al., 2021; Zhang et al., 2021). To determine if this is the case for SARS-CoV-2, we used two model systems to determine if 2-E+2-3a expression activates the NLRP3-I via the mitochondrion. First, we transformed 293T cells with plasmids encoding the components of the NLRP3-I, thus reconstituting the inflammasome (Shi et al., 2016) (**Figure 3A**). Second, we transduced THP1 cells which are a human acute monocytic leukemia derived cell line with the 2-E+2-3a expression vector. The transduced THP1 cells were then treated with phorbol ester (PMA) to generate macrophages and tthe macrophages were treated with LPS + nigericin (Pan et al., 2021) (**Figure 3B**). The expression of 2-E+2-3a in both cell systems, 293T (**Figure 3C**) and THP1 macrophages (**Figure 3D-F**), resulted in enhanced secretion of NLRP3-activated IL-1β secretion.

**Figure 3.**
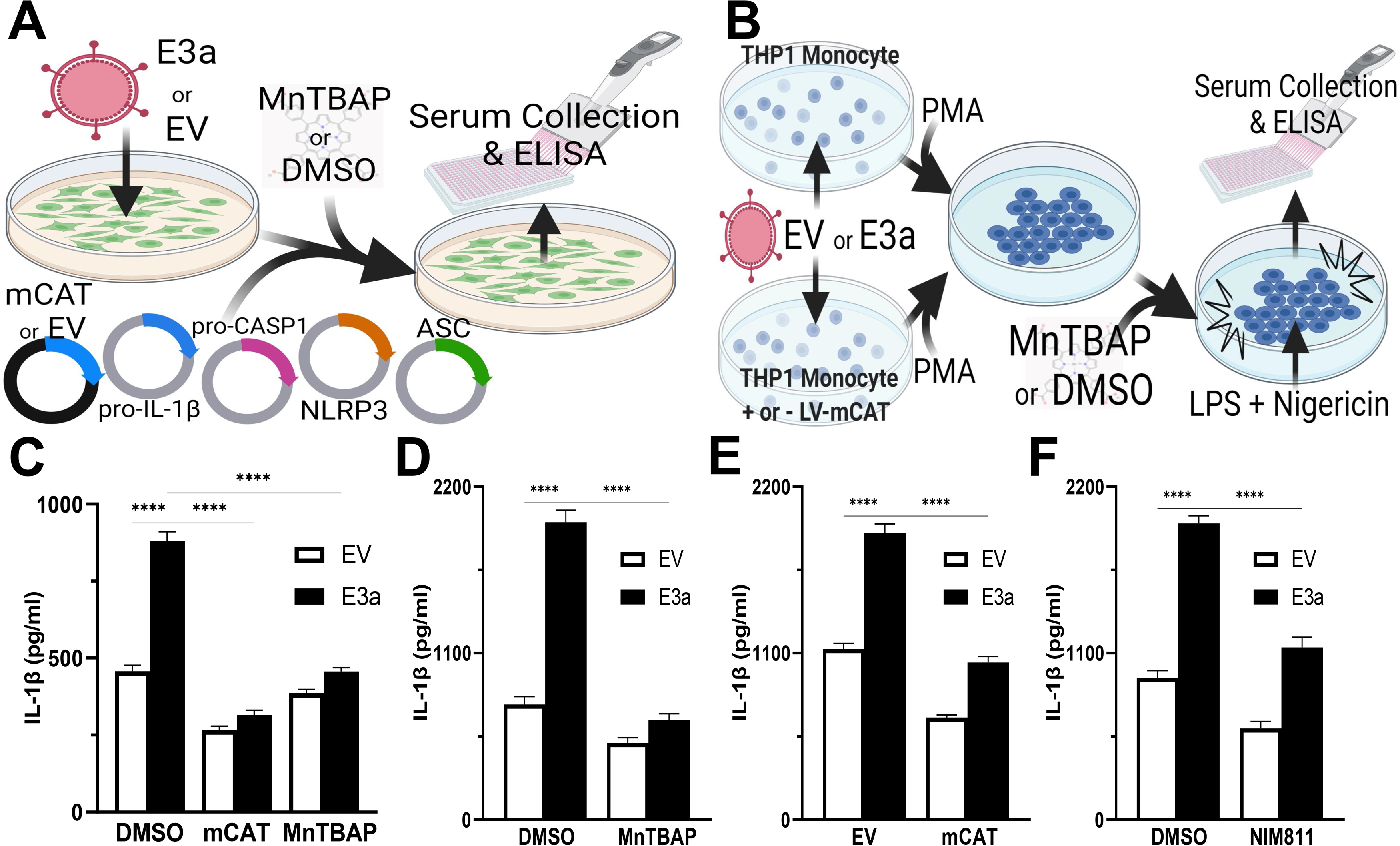
mROS and the mtPTP are required for activation of the NLRP3-I by the 2-E+2-3a viroporins. **A-B)** Experimental design used to assess NLRP3-activated by IL-1β in cell-free supernatants quantified by ELISA. **A)** 293T cells with an NLRP3-I reconstitution system (NLRP3, ASC, pro-CASP1, pro-IL-1β) and **B)** THP1 differentiated into macrophages and primed with LPS + nigericin. **C)** 293T cells transfected with LV-EV or LV-E3a were transformed the NLRP3-I plasmids and p-mCAT or its control plasmid p-EV, or cultured in the presence or absence of 50 μM MnTBAP. **D, F)** THP1 cells were transduced with LV-EV or LV-E3a, differentiated into macrophages, treated with LPS and nigericin, and treated with 100 μM MnTBAP or 10 μM NIM811, the supernatants analyzed for IL-1β by ELISA. **E)** THP1 cells stably expressing LV-mCAT or control were infected with LV-EV or LV-E3a, differentiated into macrophages, and supernatant IL-1β levels determined via ELISA. Error bars represent SEM from 3 independent experiments; statistically significant data is indicated with asterisks (*).

We then confirmed that 2-E+2-3a expression activates the NLRP3-I and IL-1β secretion via increased mROS production. 293T cells expressing the inflammasome proteins (**Figure 3C**) and LPS-nigericin treated THP1 macrophages (**Figure 3D&E**) were treated with mitochondrially targeted antioxidants, transformation with mCAT or treatment with MnTBAP. Both mCAT expression and MnTBAP treatment impaired IL-1β secretion.

We then determined if mROS activation of the NLRP3-I was mediated by release of an oxidized mitochondrial component via the mtPTP, which has been conjectured but not proven. We treated 2-E+2-3a transformed THP1 macrophages with the specific mtPTP inhibitor N-methyl-4-isoleucine-cyclosporin (NIM811). NIM811 blocks the mtPTP by binding to cyclophilin D, analogous to cyclosporin A (CsA) but without calcineurin inactivation (Dittmar et al., 2021; Tóth et al., 2019; Zhang et al., 2020). NIM811 treatment suppressed the secretion of IL-1β following LPS + nigericin activation of the THP1 macrophages demonstrating for the first time the mtPTP is the route by which an oxidized mitochondrial factor reaches the NRLP3-I (**Figure 3F**).

Thus, we have demonstrated that co-expression of 2-E+2-3a enhances Ca^++^ leakage into the cytosol, increasing levels of cytosolic and mitochondrial Ca^++^. This 2-E+2-3a mediated increase in mitochondrial Ca^++^ induces the production of mROS, which in turn activates the NLRP3-I, via mtPTP transport of an oxidized mitochondrial product, stimulating the secretion of IL-1β. Increasing mitochondrial antioxidant defenses through treatment with the pharmacological mROS scavenger MnTBAP, or genetic expression of mCAT, detoxifies 2-E+2-3a induced mROS and blocks activation of the NLRP3-I. Together these findings reveal that the mechanism by which 2-E+2-3a engage the NLRP3-I is via viroporin manipulation of mitochondrial physiology.

## DISCUSSION

Because of the importance that activation of the inflammasome by SARS-CoV-2 has on the severity of COVID19, we set out to define the physiological process by which the virus activates the NLRP3-I in hopes of identifying drug targets to mitigate the cytokine storm. We found that the viroporins 2-E+2-3a were central to the activation of the NLRP3-I and this occurred through a mitochondrial innate immunity signal transduction pathway. Expression of viroporins 2-E+2-3a results in the release of ER and extracellular Ca^++^ into the cytosol where the Ca^++^ is taken up by the mitochondrion via the mitochondrial calcium uniporter. The mitochondrial Ca^++^ activates the tricarboxylic acid cycle dehydrogenases generating excess NADH (Denton, 2009). This overloads the electron transport chain producing increased mROS. The increased mROS oxidizes a mitochondrial factor that is released through the mtPTP to bind and activate the NLRP3-I.

While our experiments, did not directly identify the mitochondrial factor released through the mtPTP, other recent studies have shown that this factor is oxidized mtDNA which is a ligand of NLRP3-I (Xian et al., 2021; Zhong et al., 2018). Thus, we complete the mitochondrial innate immune activation pathway by showing that release of Ox-mtDNA is via the mtPTP.

Demonstration that SARS-CoV-2 activated the inflammasome via the mitochondria provides new approaches to mitigating the severity of the cytokine storm. Previous studies have indicated that generalized antioxidants such as N-acetyl cysteine (Garozzo et al., 2007; Geiler et al., 2010; Ghezzi and Ungheri, 2004), glutathione (Cai et al., 2003; Nencioni et al., 2003), and catalase (Shi et al., 2014; Shi et al., 2010) can reduce viral propagation and pathology. Our data extend these observations by indicating that the therapeutic potential of drugs will be enhanced if they are mitochondrially targeted antioxidants such as MnTBAP (Melov et al., 1998; Tong et al., 2007), EUK-8 and EUK-134 (Melov et al., 2001) and or inhibitors of the mtPTP such as NIM811 (Dittmar et al., 2021; Tóth et al., 2019; Zhang et al., 2020).

### Limitations of the Study

A unique feature of this research is the discovery that SARS-CoV-2 viroporins active the inflammasome via the mitochondrion through elevated mitochondrial Ca^++^ and mROS and the mtPTP. However, we have not identified this released mitochondrial oxidized product. Rather, we relied of the publications of others implicating Ox-mtDNA.

## ACKNOWLEDGMENTS

The authors thank Dr. Bruce Beutler for gifts of expression plasmids and Dr. Cristina Mammucari (Department of Biomedical Sciences, University of Padua, 35131 Padua, Italy) and Dr. Kevin Foskett (Dept Cell and Developmental Biology, University of Pennsylvania, 19104 Philadelphia, Pennsylvania) for gifting MCUi11, which they purchased from AKos Consulting & Solutions GmbH. This work was supported by DOD grant W81XWH-21-1-0128 awarded to D.C.W.

## AUTHOR CONTRIBUTIONS

Conceptualization: J.W.G., D.C.W.; Methodology: J.W.G., P.P, D.C.W., A.A., D.M.; Literature and concept integration: J.W.G., P.P, D.C.W., A.A., D.M.; Formal Analysis: J.W.G., A.A., D.M., D.C.W.; Writing – Original Draft: J.W.G.; Investigation: J.W.G., A.A., D.M., D.C.W., T.L.; Sample Collection: J.W.G., T.L.; Writing – Review & Editing: J.W.G., D.C.W., A.A., D.M.; Visualization: J.W.G., D.C.W., A.A., D.M.; Supervision: J.W.G., P.P, D.C.W., A.A., D.M.; Funding Acquisition: D.C.W.

## MAIN TABLES AND LEGENDS

**Non-Applicable**

## STAR★METHODS

### RESOURCE AVAILABILITY

#### Lead Contact

Further information and requests for resources and reagents should be directed to and will be fulfilled by the Lead Contact, & Douglas C. Wallace.

#### Materials Availability

This study did not generate new unique reagents.

#### Data and Code Availability

This study did not generate any unique datasets. All data is included in the manuscript or supplementary file.

## EXPERIMENTAL MODEL AND SUBJECT DETAILS

### Cells, Infections, & Reagents

293T & THP1 cells were obtained from the American Type Culture Collection (ATCC). Cells were grown at 37°C with an atmosphere of 98% humidity and 5% CO2. 293T cells were maintained in Dulbecco’s modified Eagle’s medium + GlutaMAX™ supplement with pyruvate (GIBCO), 1% non-essential amino acids (SIGMA), and 10% fetal bovine serum (FBS) (Takara Bio). THP1 cells were grown in RPMI 1640 Medium (GIBCO) supplemented with 10% FBS (Takara Bio). 293T cells were infected (MOI 4) as previously described (Potluri et al., 2016). THP1 cells were infected (MOI 8) with the addition of 5 μg/ml polybrene (VectorBuilder) and spin-inoculated at 700×g for 25min.

### Plasmids, Viral Vectors, THP1 stable-transformants

#### Plasmid Vectors

To express the components of the NLRP3-inflammasome (NLFP3-I), we utilized four plasmids expressing mouse NLRP3 (pcDNA3-N-Flag-NLRP3, Addgene plasmid # 75127), ASC (pcDNA3-N-Flag-ASC1, Addgene plasmid # 75134), CASP1 (pcDNA3-N-Flag-Caspase-1, Addgene plasmid # 75128) and pro-IL-1B (pCMV-pro-Il1b-C-Flag, Addgene plasmid # 75131), The use and construction of the NLRP3-I expression plasmids were previously described (Shi et al., 2016).

The plasmid vector used to express mitochondrial-targeted catalase (mCAT) and its respective control vector were p-mCAT (VectorBuilder ID VB170403-1078nzg) and p-EV (VectorBuilder ID VB210726-1273jte), vectorbuilder.com. The p-mCAT transgene cassette is transcribed from the 212 nucleotide elongation factor α1 “short” (EFS) promoter. The EFS promoter transcribes a polycistronic transcript encoding EGFP (enhanced green fluorescent protein), a self-cleaving 2A peptide site, followed by mCAT, terminated by a simian virus 40 (SV40) late polyA sequence. p-EV is identical to the p-mCAT construct, except lacking the mCAT sequence.

#### Lentiviral Vectors

The lentiviral vector used to co-express 2-E+2-3a was LV-E3a (Vectorbuilder ID VB210112-1153ufz) and its respective control vector LV-EV (Vectorbuilder ID VB210112-1153ufz). The LV-E3a vector contains the *cytomegalovirus* (CMV) promoter, the 2-E+2-3a viroporins obtained from Gordon *et al*. 2020 (Gordon et al., 2020b) separated by a 2A peptide site, and terminated by a SV40 late polyA sequence cloned into the LV-EV vector. The viroporins were modified by addindg anATG codon 5’ and three N-terminal FLAG-tags were added to the 3’ end of each viral protein, and transcribed from the the proteins. LV-EV is an empty vector.

The lentiviral vector expressing our mCAT and its respective control vector, LV-mCAT (VB210909-1242kdf) and LV-EV(mCAT) (VB900122-0484ubz) were constructed and packaged by VectorBuilder. The LV-mCAT vector includes the EFS promoter, EGFP, 2A peptide site, mCAT,, and SV40 late polyA sequence. The LV-EV(mCAT) control vector lacks EGFP and mCAT.

#### THP1 mCAT stable-transformants

THP1 cells were transduced with LV-mCAT or empty vector and selected with puromycin. Expression of mCAT was \validated stable-transformants by EGFP fluorescence.

## METHOD DETAILS

### Cell Staining

293T cells were plated at a density of 45 × 10^3^ in 96-well 0.2% gelatin-coated (ScienCell) glass-bottom plates with high-performance #1.5 mm cover glass (Cellvis). Twenty-four hours post-plating, sub-confluent monolayers of 293T cells were transduced with LV-EV or LV-E3a. Twenty-four hours post-transduction, cells were washed two times with phosphate-buffered saline (PBS), then stained. For determination of mROS levels, cells were co-stained with 3 μM MitoSOX™ Red (MitoSOX, mitochondrial superoxide indicator) and 50 nM MitoTracker™ Deep Red FM (MTDR) for 30 min at 37°C. For assaying mROS levels after treatment with Thapsigargin (TG) using the plate reader, cells were stained with 3 μM MitoSOX for 30 min at 37°C.To quantify mROS after staining, cells were washed three times in PBS, maintained in FluoroBrite™ DMEM (GIBCO) supplemented with 12.5 mM HEPES (SIGMA) and 1% non-essential amino acids (SIGMA), and the fluorescence measures

To determine cytosolic Ca^++^ levels, cells were washed three times with Tyrode’s Salts (Sigma-Aldrich), stained for 40 min with 2 μM Fura Red™, acetoxymethyl ester (AM), cell-permeant (Fura-Red) in 0.02% pluronic F127 (Pluronic® F-127) detergent. To determine mitochondrial Ca^++^ levels, cells were washed three times with Tyrode’s Salts, stained for 40 min with 10 μM Rhod-2, AM, cell-permeant (Rhod2) and 50 nM MTDR in 0.02% pluronic F127. After staining cells were washed three times and maintained in Tyrode’s Salts and florescence measured. To determine mitochondrial Ca^++^ levels after treating with TG, cells were stained for 40 min with 10 μM Rhod2, washed three times and maintained in Tyrode’s Salts, and florescence measured.

### SpectraMax Plate Reader Assay

#### Measurement of mROS and cytosolic and mitochondrial Ca^++^ levels

After staining cells (see “Cell Staining”), mean fluorescence was assessed using the *SpectraMax® Paradigm® Multi-mode Detection Platform, equipped with a* Tunable Wavelength (TUNE) Detection Cartridge (Molecular Devices). was quantified by MitoSOX fluorescence (ex:540 nm, em:590 nm) and MTDR (ex:633 nm, em:680 nm) and MitoSOX/MTDR calculated. Rhod2 fluorescence for mitochondrial Ca^*++*^ level (ex:540 nm, em:590 nm) and MTDR (ex:633 nm, em:680 nm), and Rhod2/MTDR calculated. Fura-Red fluorescence for cytosolic bound-Ca^++^ level (ex:405 nm, em:637 nm) and unbound-Ca^*++*^ (ex:514 nm, em:672 nm) state. The ratio of bound-Ca^++^/unbound-Ca^++^ was calculated.

#### Measurement of mitochondrial Ca^++^ and mROS after treating with TG

After staining cells with Rhod2 or MitoSOX (“Cell Staining”), cells were treated for 10 min with or without 10 μM mitochondria channel uniporter inhibitor 11 (MCUi11), Dimethylsulfoxide (DMSO) as a negative control. Mitochondrial Ca^++^ determined from Rhod2 (ex:540 nm, em:590 nm) and mROS from MitoSOX (ex:540 nm, em:590 nm). Cells were tthen treated with 2.5 μM TG and changes in mitochondrial Ca^++^ or mROS recorded every 15 seconds for 180 seconds. Relative change in mitochondrial Ca^++^ was calculated by dividing the average change in Rhod2 fluorescence after treatment with TG by Rhod2 fluorescence before treatment with TG. Relative change in mROS levels was calculated by dividing the average change in MitoSOX fluorescence after treatment with TG by MitoSOX fluorescence before treatment with TG.

### Confocal Microscopy

Live-cell imaging was performed using a Zeiss 710 LSM confocal microscope with an environmental chamber maintained at 37°C and 5% CO2. Laser lines used: diode 405 nm, Argon 514 nm, HeNe lasers 543 nm and 633 nm excitation wavelengths. Fluorescence quantified using the Zeiss 710 LSM confocal microscope was analyzed using ImageJ. Fluorescence intensities of stained cells were normalized to the unstained negative cells. A minimum of 60 images was taken for each condition across at least three independent experiments.

### Catalase Assay

Catalase activity was assessed through cleavage of H2O2 in 293T cell lysates collected twenty-four hours post-transfection with p-mCAT or p-EV, using a Catalysis Activity Kit (Abcam, ab83464).

### Western blot analysis

Twenty-four hours post-transduction, cells were washed once with cold PBS and then lysed with 2.5% n-Dodecyl-B-D-Maltoside in 20 mM HEPES (pH 7.4), 50 mM βglycerophosphate, 2 mM EGTA, 10% (v/v) glycerol, and 0.01% Bromophenol blue. Lysates were electrophoresed on 4 to 12%, Bis-Tris gels (Invitrogen) SDS-polyacrylamide NuPAGE™ gels Gels were ransferred to a nitrocellulose membrane by the iBlot Gel Transfer System (Invitrogen), membranes blocked for one hour in 5% nonfat milk in 25 mM Tris-HCl, 150 mM NaCl, 0.1% Tween 20 (TBST buffer and incubated overnight at 4°C with shaking in primary antibody diluted 1:1000 in TBST. Membranes were then washed three times with TBST and incubated with Alexa Fluor-conjugated secondary antibodies for one hour at room temperature. Protein levels were quantified using the Odyssey imaging system (LiCOR Biosciences) using. GAPDH as a loading control.

### Detection of secreted IL-1β in 293T cells with a reconstituted NLRP3-I and THP1 macrophages

293T cells were plated at a density of 100 × 10^3^ in 24-well plates. Twenty-four hours post-plating sub-confluent monolayers of 293T cells were infected with LV-EV or LV-E3a. Six hours post-infection cells were co-transfected using the TransIT-X2® Dynamic Delivery System (Mirus Bio) with the plasmids encoding the components of the NLRP3-I (Shi et al., 2016), followed by p-mCAT or p-EV transduction. Twenty-four hours post-transfection, cells were washed two times with PBS, then cultured with or without the addition of 50 uM MnTBAP. Cell lysates and culture supernatants were collected twelve hours post-treatment andcentrifuged to remove cell debris. Supernatant IL-1β was quantified by ELISA (Abcam, ab197742).

THP-1 cells were plated at a density of 100 × 10^3^ in 96-well plates. THP1 cells were infected with LV-EV or LV-E3a. Six hours post-infection, THP1 cells were differentiated into macrophages with 50 ng/ml Phorbol 12-myristate 13-acetate (PMA) overnight. After differentiation, cells were washed two times with PBS, and fresh media was added with addition of 100 μM MnTBAP or 10 μM N-methyl-4-isoleucine-cyclosporin (NIM811). Nine hours post-treatment with MnTBAP or one hour post-treatment with NIM811 THP1 macrophages were stimulated with 100 ng/ml Lipopolysaccharides (LPS) and 2.5 mM of nigericin for nine hrousSupernatants were collected, centrifuged, and the amount of supernatants IL-1β measured by ELISA (Abcam, ab46052). THP1 cells stably expressing LV-mCAT or control vector, were infected with LV-EV or LV-E3a, and 6 hours post-infection the THP1 cells were differentiated into macrophages, and supernatant IL-1β quantified via ELISA.

## QUANTIFICATION AND STATISTICAL ANALYSIS

### Statistical analysis

One-way ANOVA was performed for statistical differences between three or more groups, followed by a post hoc Tukey’s HSD test to test for statistical differences. For studies that require a quantitative evaluation between two groups, statistical significance was determined using unpaired two-tail student’s t-test. All data are reported as mean ± standard error of the mean (SEM). All statistical analysis was done on GraphPad Prism 9.01. For student’s t-test * = p < value 0.05, ** = p < value 0.01, *** = p < value 0.001, **** = p < value 0.0001).

## SUPPLEMENTAL VIDEO, DATA, AND EXCEL TABLE TITLE AND LEGENDS

**Non-Applicable**

---

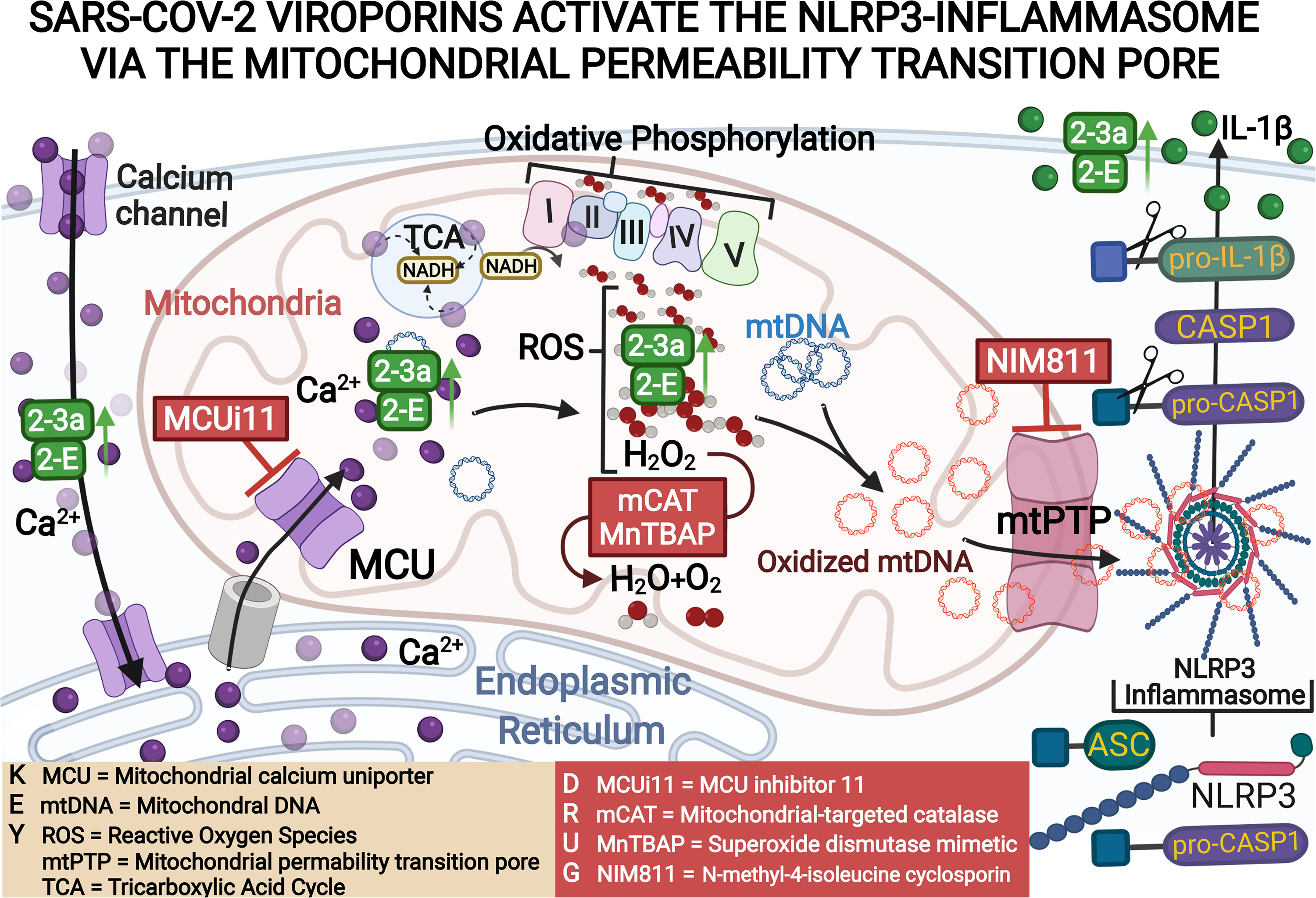

